# Improved efficacy and tolerability of antisense oligonucleotide with Guanidine- bridged nucleic acid

**DOI:** 10.64898/2026.01.08.698310

**Authors:** Takaaki Kawanobe, Ajaya Ram Shrestha, Hideaki Tomita

## Abstract

Guanidine- bridged nucleic acid (GuNA) is a bridged nucleic acid analog with high binding affinity towards complementary strands along with high nuclease resistance. GuNA has been developed to improve pharmacokinetics and safety profiles of phosphorothioate modified gapmers. Here, we evaluated antisense oligonucleotides (ASOs) modified with combination of GuNA and 2’-O-methoxyethyl (MOE) could significantly improve KD activity in vitro and in vivo. Long-term efficacy evaluation showed that intracerebroventricularly administered GuNA modified gapmers stayed active for over 24 weeks in mouse brain. Furthermore, we found that GuNA-modified gapmers could evade lymphocyte-derived immune responses and kept ASO-induced toxicity in check. Taken together, the results of this study demonstrated that GuNA modification can improve the potential of ASOs, especially in the central nervous system.

## Introduction

Antisense oligonucleotides (ASOs) are short, synthetic, single-stranded deoxyribonucleotides regulating the target RNA expression through Watson-Crick base pairing. A gapmer-type ASO (gapmer) consists of a central region of deoxyribonucleotides (gap) flanked by 2′-modified sugar nucleotides region (wings), which can induce RNase H1-mediated target degradation. Among the 14 clinically approved ASOs to date, 7 are gapmer ASOs^1^. In clinic, the approved gapmers are given to patients once a week subcutaneously or once a month intrathecally for maximal effects^1,2^. One of the reasons for the high frequency of dosing is because of susceptibility of the oligonucleotides towards degradation by endogenous intracellular nucleases^3^. Therefore, high nuclease resistance is crucial to develop an ideal long-effective therapeutic ASO. Many chemical modifications are in practice to improve biostability and other properties of an oligonucleotide^4^. The internucleotide linkages of an ASO are modified with phosphorothioate (PS) bonds, where the non-bridging oxygen is replaced with a sulphur. The modification can improve nuclease stability, pharmacokinetics, and cellular/nuclear uptake, which makes it a favourable and unavoidable backbone modification for therapeutic ASOs^5^. Modifications of the sugar moiety, such as 2’,4’-bridged nucleic acid/locked nucleic acid (2’,4’-BNA/LNA), provide a gapmers strong affinity for target RNA and high nuclease resistance^6,7^. Furthermore, some 2’,4’-BNA/LNA analogs can reduce ASO-mediated hepatotoxicity and alter biodistribution of ASOs^8,9^. Therefore, sugar modification of ASOs is an effective strategy to enhance potency and metabolic stability^10^. Although LNA could show promising properties, there has not been any FDA-approved ASO therapeutics with the modification because of the safety concern of LNA modified oligonucleotides in animal studies^11,12^. Currently, 2’-O-methoxyethyl (MOE) modification has been majorly used in clinical ASO candidates; most of the approved ASOs are modified with the modification. However, inflammatory adverse events (AEs) such as flu-like symptoms have been reported even with the approved ASOs with MOE modification^13,14^. Therefore, it is crucial to develop in vitro tests for chemically modified ASOs that can predict ASO-induced inflammatory responses at an early stage before development^15–17^. The ASO-mediated immune response is due to reactivity to toll-like receptor 9 (TLR9). TLR9 reactivity can be evaluated by using human PBMC assays. However, donor variability makes it difficult to obtain clear and consistent results. Recently, B cells from PBMC can predict ASO-induced inflammatory responses like PMBC, which further specified that ASOs trigger B cells activation via TLR9 and led to cytokine release^15^.

Here, we report the reduction of immunotoxicity without compromising potency by the incorporation of GuNA into wing regions of MOE modified gapmers, including ISIS 301012 which has been clinically observed to cause injection site inflammatory reactions^13^. These MOE gapmers are highly effective and GuNA modification further improved their safety for systemic and central nervous system application. Furthermore, long-term durability evaluation in mouse brain demonstrated potential for improved stability of GuNA modified gapmers over gapmers modified only with MOE. Overall, our results showed that GuNA incorporated MOE gapmers could be a new scaffold to enhance therapeutic profiles of the ASOs and bring findings to avoid of modification-derived immunotoxicity.

## Materials and Methods

### Oligonucleotide synthesis and purification

Antisense oligonucleotides were designed with MOE, GuNA and LNA as shown in Fig.1A, 1B and supplementary Fig.1A. These gapmers were synthesized on a 10 μmol scale by using an ns-8II Synthesis System (Genedesign, Inc.) according to the standard phosphoramidite protocol. 5-[3,5-bis(trifluoromethyl)phenyl]-1*H*-tetrazole (Activator 42®) was used as the activator, and Cap mix A (10% acetic acid in tetrahydrofuran) and Cap mix B (10% 1-methylimidazole in tetrahydrofuran/pyridine) were used as the capping agents. 0.05 M DDTT SulfurizingReagent(3-((dimethylaminomethyliden)amino)-3*H*-1,2,4-dithiazol-3-thione) (ChemGenes) in pyridine/acetonitrile was used for thiolation and 0.02 M iodine in tetrahydrofuran/water/pyridine was used for oxidation. The standard synthesis cycle (trityl off mode) was used for assembly of the reagents and synthesis of the oligonucleotides, except that the coupling time was extended to 30 min for the GuNA, MOE monomers. The phosphoramidites were dissolved in anhydrous acetonitrile at 0.1 M concentration. Unylinker CPG-solid supports from Chemgenes were used. After synthesis, the synthesized oligonucleotides were treated with 20% diethylamine in acetonitrile for 1h and cleaved from the solid support and deprotected by treatment with 28% aq. ammonia at 55 ° C for 8 h. The oligonucleotides were purified by reverse-phase chromatography on Agilent HPLC system using mobile phase 100 mM HFIP/8.6 mM TEA in water (mobile phase A) or methanol (mobile phase B), following by ion-exchange chromatography on a Akta pure 25 using 0.01M phosphate buffer (pH 8.0) and 2M NaBr. Pure fractions were desalted on sephadex G-25 column (cytiva Hiprep 26/10 desalting) and lyophilized. Purity and mass of oligonucleotides were determined by using ion-pair LCMS.

**Fig. 1.**
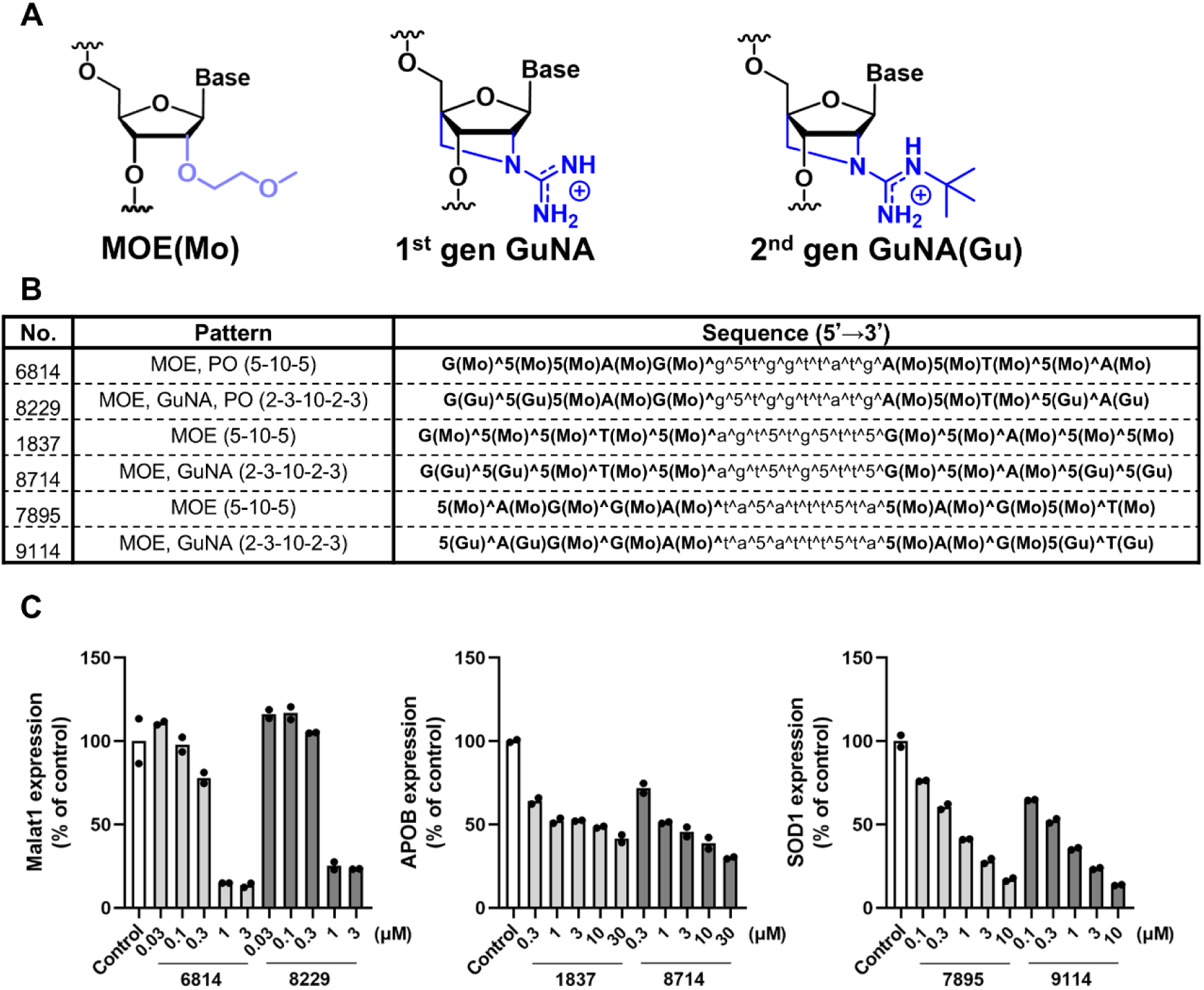
(A) Chemical structure of modified nucleotide. (B) Chemical modifications and sequences of modified gapmer were listed. The numbers in parentheses indicate the number of modified nucleoside in the order of 5’wing-gap-3’wing. *Gu* indicate GuNA, *Mo* MOE, *^* phosphorothioate bond, *5* 5’-methylcytosine, *Upper letter* modified nucleoside and *lower letter* DNA. Bond between nucleoside without *^* is phosphodiester bond. (C) Percentage of mRNA expression relative to untreated control. Cells were treated with modified gapmer by gymnotic uptake for 72 hours. (biological replicate, n=2)

### Cell lines

HuH-7 cells were purchased from JCRB Cell Bank (Osaka, Japan). KELLY cells and Neuro-2a cells were purchased from ECACC. BJAB cells were purchased from the Leibniz Institute DSMZ (Brunswick, Germany). BJAB cells were cultured in growth medium supplemented with 20% fetal bovine serum (FBS; Gibco/Thermo Fisher Scientific). The other cells were cultured in growth medium supplemented with 10% FBS.

### Modified gapmer knockdown experiment

HuH-7 cells, KELLY cells and Neuro-2a cells split into 96-well plates were cultured in a medium. HuH-7 and KELLY cells were gymnotically transfected with modified gapmer and incubated for 72 hours. Neuro-2a cells were transfected by using lipofectamine 3000 and incubated for 24 hours. The cells were stored at −70 °C until processing for RNA extraction.

### Human PBMC assay

Donor-derived human PBMCs were purchased from Lonza (CC-2702P; Walkersville, USA). The cells were resuspended in RPMI1640 supplemented with 10%FBS. The cells were plated at 2.5×10^5^ in 100 µL/well of 96-well plate. 25 µL/well of 5× concentrated modified gapmer diluted in RPMI1640+10% FBS was added. The plated cells were incubated for 24 hours at 37 °C; 5% CO_2_. Plates were centrifuged at 510 *g* for 10 min before sampling supernatants. Cell supernatants were stored at −70 °C until processing for IL-6 ELISA. Detection of IL-6 release from cells were according to manufacturer instructions using ELISA MAX™ Deluxe Set Human IL-6 (BioLegend).

### BJAB assay

For BJAB assay, BJAB cells were washed in serum free RPMI1640 and resuspended in serum free RPMI1640. The cells were plated at 2×10^5^ in 50 µL/well of 96-well plate. 50 µL/well of 5× concentrated modified gapmer diluted in serum free RPMI1640 was added. The plated cells were incubated for 24 hours at 37 °C; 5% CO_2_. Plates were centrifuged at 510g for 10 minutes and stored at −70°C until RNA extraction.

### In vivo studies in mice

Animal experiments were conducted in accordance with a protocol approved by the Animal Care and Use Committee at Luxna Biotech. Female 5 weeks old mice (ICR) were purchased from CLEA Japan and allowed to acclimatize for 1 week before the experiments. Modified gapmers were administered via intracerebroventricular (icv) single injection at a dose of 20 nmol/head or indicated dose. Tissues collected 2, 8, 16 and 24 weeks after administration. Laparotomy was performed for all animals under isoflurane anesthesia (1.5-3%). After macroscopic examination for abdominal organs and tissues, animals were euthanized by exsanguination, and the brain compartments were dissected. Tissue samples were subsequently stored in RNAlater Stabilization Solution (Thermo Fisher Scientific) for RNA expression analysis. For immunohistochemistry the whole brains were obtained 2 weeks after administration, immersed and fixed in 10 vol% neutral buffered formalin at room temperature.

### RNA extraction and qPCR analysis

RNA extraction from cells was according to manufacturer instructions using KingFisher Flex (Thermo Fisher Scientific) in the MagMAX™ mirVana™ RNA Isolation Kit (Thermo Fisher Scientific). RNA extraction from tissues was according to manufacturer instructions following beads homogenization using Tissue Lyser II (Qiagen) in TriPure isolation reagent (Roche). TaqMan qRT–PCR was performed using QuantiNova PCR Kits (Qiagen) and TaqMan Gene Expression Assays (Thermo Fisher Scientific). In brief, reverse transcription was performed at 50 °C for 10min; then the reactions were denatured at 95 °C for 5min and 40 cycles of PCR reactions were performed at 95 °C for 15 s and 60 °C for 30 s by using CFX Connect Real-Time PCR detection system (Bio-rad). The TaqMan probe product IDs were human ACTB (Hs99999903_m1), human APOB (Hs00181142_m1), human SOD1 (Hs00533490_m1), human CCL22 (Hs01574247_m1), mouse Gapdh (Mm99999915_g1), mouse Malat1 (Mm01227912_s1), and mouse Cd68 (Mm03047343_m1). RNA expression levels were analysed by CFX Maestro Software (Bio-Rad).

### Immunohistochemistry for modified gapmer

Immunostaining studies were performed at Axcelead Drug Discovery Partners, Inc. (Kanagawa, Japan). The paraffin embedded whole brain tissues were sectioned. Modified gapmer was detected with anti-PS antibody (Axcelead DPP), and cell-specifc markers were detected with NeuN (abcam), GFAP (Dako), Iba-1 (abcam), TPPP (abcam), and CD31(Cell Signaling Technology: CST) antibodies.

## Results

### GuNA modified gapmers were as potent as other modified gapmers in RNA knockdown

The knockdown (KD) activity of 1^st^ generation GuNA-modified gapmers has already been reported^9,18^. Here, we tested the KD activity of modified gapmer by the combination of 2^nd^ generation GuNA, i.e. GuNA[*t-*Bu] (hereafter referred to as GuNA) and MOE (Fig. 1A)^19^. We used the sequences of ISIS 301012, ISIS 333611, ISIS 626112 for the modification. The modified gapmers were designed substituting two MOEs with two GuNAs at both ends of each wing (Fig.1B). All modified gapmers were tested in cultured cells to evaluate KD activity. In all cases, the target gene expressions were similarly suppressed with GuNA modified gapmer (LX-A8229, LX-A8714 and LX-A9114) and MOE modified gapmer (LX-A6814, LX-A1837 and LX-A7895) (Fig.1C). In addition to LX-A6814, we also designed a modified gapmer based on LX-A7786 targeting the Malat1 gene (supplementary Fig.1A). LX-A7787 and LX-A7788 modified with LNA or GuNA respectively showed KD activity equivalent to that of MOE (supplementary Fig.1B). These results suggested that KD activity of an ASO is not compromised by the introduction of GuNA in the MOE-modified sequence.

### GuNA modification can mitigate immunotoxicity

ASO-induced immunotoxicity is often an issue. Several approaches have been reported to avoid the toxicity, such as incorporation of sugar modification in the ASO^15,20,21^. To test whether GuNA modification can reduce immunotoxicity, we replaced MOE to GuNA in sequences known to be immunotoxic. We measured the amount of IL-6 released from human PBMCs to evaluate immune reaction of MOE- and GuNA-modified gapmers. Interestingly, all of the GuNA-modified gapmers (LX-A8229, LX-A8714 and LX-A9114) significantly supressed release of IL6, whereas all of the MOE modified gapmers showed significant release of IL6 (LX-A6814, LX-A1837 and LX-A7895) in PBMC (Fig.2A). To verify this immunosuppressive effect of GuNA modification, we used BJAB cells, a human B-cell lymphoma cell line. It has been reported that TLR9 activation on B-cells is known to mediate ASO-induced immunotoxicity^22^. Furthermore, the immune reactivity of BJAB cells shows better correlation to clinical adverse events. As seen in PBMC, GuNA-modified gapmers (LX-A8229, LX-A8714 and LX-A9114) significantly suppressed the expression of CCL22, whereas MOE-modified gapmers (LX-A6814, LX-A1837 and LX-A7895) induced strong CCL22 expression (Fig.2B). These results verified the mitigating effects on immunotoxicity by GuNA modification in PMBC and strongly suggested that GuNA modification can be highly effective to negate immunotoxicity in general.

**Fig. 2.**
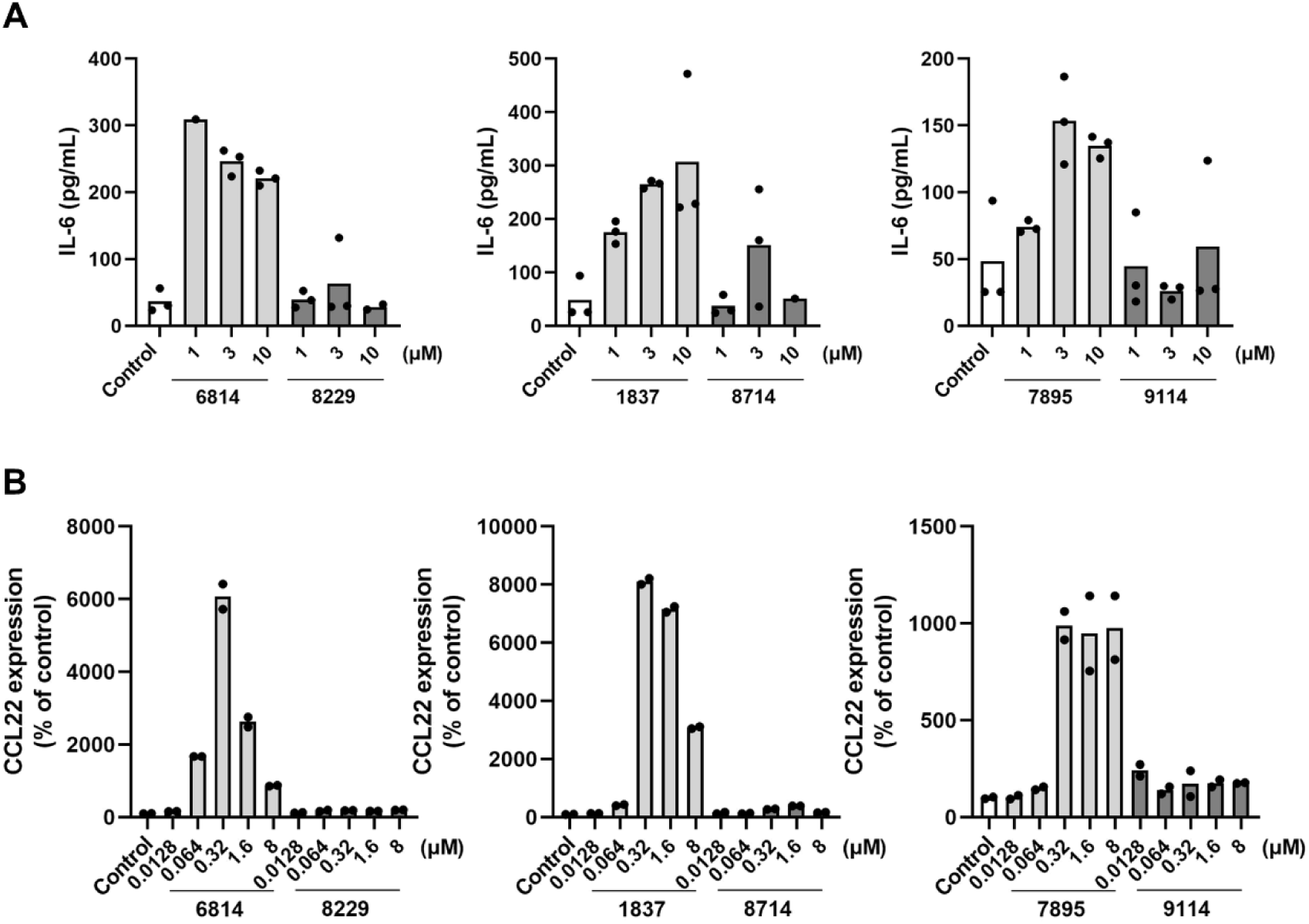
GuNA suppressed immunotoxicity. (A) Amount of IL-6 release relative to untreated control. Cells were treated with modified gapmer by gymnotic uptake for 24 hours. (n=3) (B) Percentage of CCL22 mRNA expression relative to untreated control. Cells were treated with modified gapmer by gymnotic uptake for 24 hours. (n=2)

### GuNA can improve durability and safety of modified gapmers in vivo

Next, we used modified gapmers targeting the Malat1 gene to evaluate the duration and distribution of the modified gapmers in the mouse brain after icv administration. The distribution of MOE-modified gapmers in the brain after icv administration in mice has already been reported^24^. We investigated the differences between MOE (LX-A6814) and GuNA (LX-A8229) modified gapmers. All modified gapmers were tested in ICR mice after single-dose icv administrations to evaluate modified gapmer potency for 24 weeks (Fig.3A). Malat1 gene suppression was detected in the cerebral cortex, brainstem, cerebellum, and spinal cord by both LX-A6814 and LX-A8229 for over 24 weeks. While MOE-modified gapmer (LX-A6814) showed reduction of KD activity from 16 weeks onward, GuNA-modified gapmer (LX-A8229) showed sustained KD activity (Fig.3B). KD activity was strongest in the brainstem and weakest in the cerebellum, which was not affected by the difference in modification between MOE and GuNA. In addition to MOE-modified gapmer (LX-A6814), we also evaluated another MOE-modified gapmer (LX-A7786) targeting the Malat1 gene (supplementary Fig.1A). When MOE was replaced with GuNA in LX-A7786, , this GuNA-modifed gapmer (LX-A 7788)also showed stronger KD activity in each brain compartment than LX-A7786 (supplementary Fig.1C, 1D). Interestingly, when MOE was replaced with LNA, another kind of the bridged nucleic acid, in LX-A7786, this LNA-modified gapmer (LX-A7787) did not improve KD activity compared to the orignial MOE-modified gapmer, LX-A7786.

**Fig. 3.**
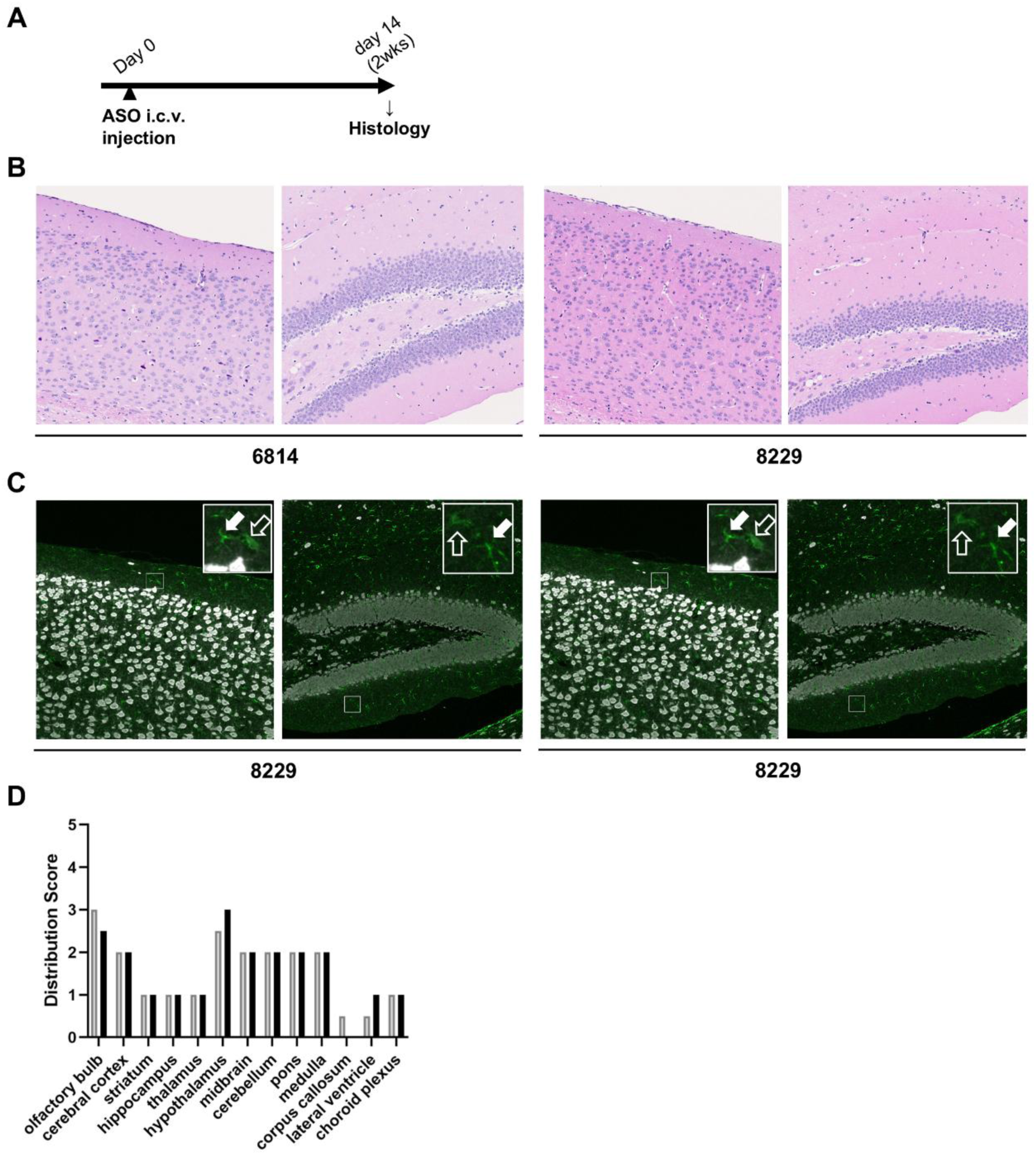
(A) Schematic representation of in vivo study design and timeline. Mice were intracerebroventricular administered a single of modified gapmers at 20 nmol/head for 2, 8, 16 and 24 weeks. (B) Time course of percentage of Malat1 mRNA expression in brain compartments (cortex, brainstem, cerebellum and spinal cord) of mice relative to saline control. (n=4) (C) Time course of tissue concentration of modified gapmer in brain compartments (cortex, brainstem, cerebellum and spinal cord) of mice. (n=4) (D) Time course of percentage of Cd68 mRNA expression in brain compartments (cortex, brainstem, cerebellum and spinal cord) of mice relative to saline control.

To examine the correlation between KD activity and the amount of remaining LX-A6814 and LX-A8229, the amount of modified gapmers distributed in the tissue was detected by LC-MS. Interestingly, although not reflected by the expression level of the Malat1 gene, the tissue concentration of LX-A8229 remained very high for a long term (Fig.3C). Mice treated with LX-A6814 exhibited late-onset neurotoxic behaviors such as hind-leg drugging, circling (data not shown) over time. It has been reported that ASO administration activates the immune system^25^, therefore we investigated long-term changes of the inflammatory marker Cd68 gene. The interferon signal-related Oasl2 gene, which was described in the paper mentioned above, did not show any significant changes (data not shown). Two weeks after the administration, MOE-modified gapmer, LX-A6814 significantly increased Cd68 gene expression and recovered over time in the brainstem and cerebellum, , whereas GuNA-modifed gapmer, LX-A8229 did not change Cd68 expression over time (Fig.3D). These results suggest that GuNA enhances the structural stability of gapmers and contributes to improved safety in-vivo.

### GuNA modification shows brain distribution similar to MOE-modification

To further characterize GuNA-modified gapmers, we investigated tissue distribution in the brain. MOE-modified gapmer (LX-A6814) and GuNA-modified gapmer (LX-A8229) were intracerebroventricularly injected into mice and the brains were examined two weeks after the the injections (Fig.4A). The histology revealed that neither LX-A6814 or LX-A8229 caused pathological abnormalities in brain regions, including the cerebrum, midbrain, and cerebellum (Fig.4B). To examine the tissue distribution of LX-A6814 and LX-A8229 in each brain region, these modified gapmers were detected by using an anti-phosphorothioate (PS) antibody. Both LX-A6814 and LX-A8229 were fairly distributed in a whole brain however both modified gapmers were relatively concentrated in the brain regions such as olfactory bulb, the hypothalamus and midbrain (Fig.4C, 4B). Furthermore, we examined if these modified gapmers show different distribution in cell types of brain. Both MOE-modified gapmer (LX-A6814) and GuNA modified gapmer (LX-A8229) were found in neurons, astrocytes, and microglia in each region of brain. GuNA modified gapmers distributed and localized similarly to MOE modified gapmers in brain. Taken together, these results suggest that GuNA modified gapmers can act in a wide range of tissues and behave similarly to MOE modified gapmers.

**Fig. 4.**
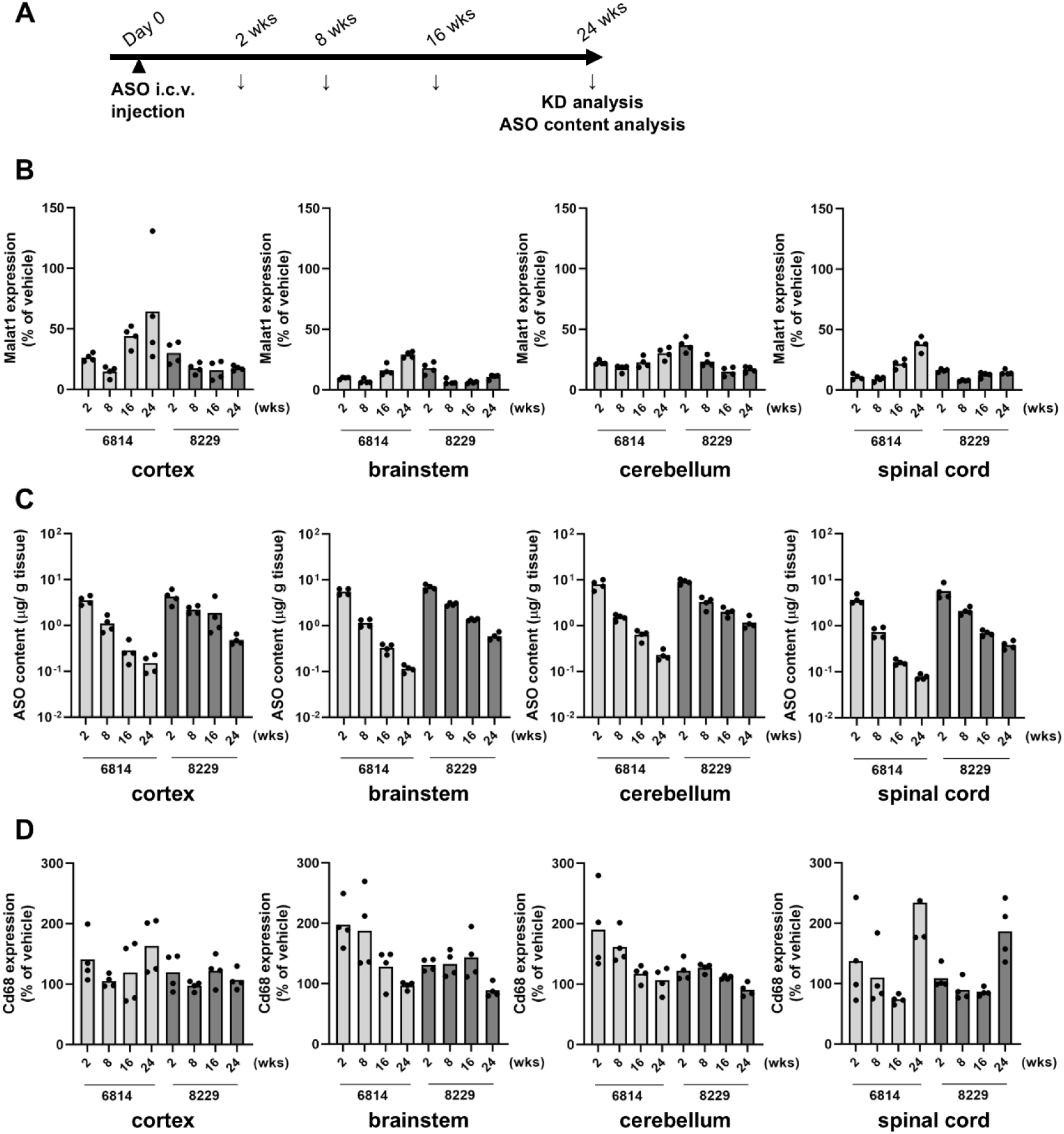
(A) Schematic representation of in vivo study design and timeline. Mice were intracerebroventricular administered a single of modified gapmers at 20 nmol/head for 2 weeks. (B) Hematoxylin and eosin staining sections of the brain from mice treated with modified gapmers or PBS and sacrificed on 2 weeks. Stained images of the cerebral cortex and hippocampus regions were shown. (C) Immunohistochemistry for modified gapmer in the centrilobular area of brain from mice treated with modified gapmers or PBS and sacrificed on 2 weeks. Modified gapmers are shown in green, and neurons in white. The inset image shows astrocytes stained with microglia.

## Discussion

It is crucial and rather difficult to maintain drug efficacy and keep toxicity in check for the development of ASO therapeutics. Modified nucleic acids, such as MOE and LNA, have been strategically used to achieve these goals, especially to reduce toxicity of ASO. In this study, we demonstrated that GuNA modified gapmers avoid typical nucleic acid-induced immune responses, such as IL-6 release and immune response gene induction in lymphocytes and evade immunotoxicity in the central nervous system. ASO-induced toxicity is due to activation of TLR9 mediated immune response. Because GuNA has bulky modifications at the 2’ position of the ribose sugar moiety^15,21^ and a basic, cationic guanidinium group with a hydrophobic tertiary butyl group, which neutralizes the charge of entire oligonucleotide (data not shown), these properties could potentially inhibit interaction of ASO (i.e., gapmers) to TLR9, leading to suppression of immunotoxicity. Furthermore, it has been reported that TLR9 dimerization followed by the recruitment of MyD88 is required for inflammatory signaling^26^. During this process, the pH of the endosomal lumen becomes acidic^27^. Because of the alkaline nature and structural bulkiness of GuNA, this potential pH resistance may inhibit specifically binding to TLR9 and lead to the suppression of immunotoxicity. It has been reported that TLR9 activation can be suppressed by bafilomycin A1 and other compounds that resist the pH decrease in the endosomal lumen^28^. GuNA may also act by similar mechanism, but further investigation is required.

Modification of sugar moieties can significantly improve stability of oligonucleotides, which can lead to long term presence and activity in tissues. GuNA modification is known to have excellent resistance against 3’-exonuclease^18,19^. It has been demonstrated that MOE-modified and GuNA-modified gapmers stayed active over 24 weeks after intravenously injected to mice. Interestingly, GuNA-modified gampers reportedly showed different biodistribution than MOE-modified gampers in mice, which is potentially due to the cationic moiety of GuNA modification^9^ Therefore, we expected that GuNA modified gapmers might show a distinctive distribution in the brain when ICV injected. However, the GuNA modified gapmers showed a tissue distribution similar to that of MOE modified gapmers.

Although GuNA modification could show the difference in bio-distribution, the difference in distribution may be subtle in brain for a few potential reasons. Transfection efficiency of modified gapmers is known to vary in different cell types^29^. In brain, modified gapmers are generally taken up to neurons inefficiently whereas they are efficiently and readily taken up to glial cells. Furthermore, unlike other organs with multiple cell types such as liver and kidney, brain consists of two major cell types, neurons and glial cells. In terms of the population of each cell type, glial cells are roughly twice as many as neurons in most of brain regions except the cerebellum^30^. When a majority of modified gapmers are readily taken up by glial cells, potential differences in distribution needed to be significant among glial cells. Therefore, it would be difficult to identify with confidence if the potential difference is present but subtle. GuNA modified gapmers maintain relatively high concentration and KD activity over time in all the brain regions except the cerebellum. When GuNA-modified gapmers were abundantly concentrated in cerebellum, KD activity of GuNA modified gapmers was relatively weaker than the other brain regions such as the cerebral cortex. It has been suggested that KD potency of modified gapmers can generally vary in different cell types. Reportedly modified gapmers are less active in granule cells than the other cell types in cerebellum^24^. This may explain the weaker KD activity of GuNA modified gapmers even with the abundant concentration in cerebellum.

Drugs for CNS diseases and disorders require long-term efficacy and safety. Intrathecal (IT) administration is generally the route of choice for central nervous system diseases. Considering IT administration is highly invasive for patients, it is preferable to treat patients with less frequent administration. GuNA-modified gapmers can sustain the availability with a long-lasting efficacy and avoid inflammatory immune responses in-vitro and in-vivo, making them highly compatible with CNS therapeutics. This study evaluated ICV administration in rodents, whereas intrathecal administration is the most commonly used route of administration for modified gapmers in clinical practice. Previous studies have reported that there is no significant difference in the pharmacokinetics of modified gapmers between ICV and IT administration^24,31^ Taken together GuNA modification can potentially overcome ASO-related issues and lead to develop ASO therapeutics with better potency and safety.

## Acknowledgement

We thank Mari Ito, Satoko Sakai, Yukai Okamoto, and Kana Sonoda of Luxna Biotech for their technical assistance. We also thank Axcelead Drug Discovery Partners, Inc. for their technical assistance in conducting Immunohistochemistry.

## Data Availability

Research data are available upon request.

---

**Supplemental Fig.1.**
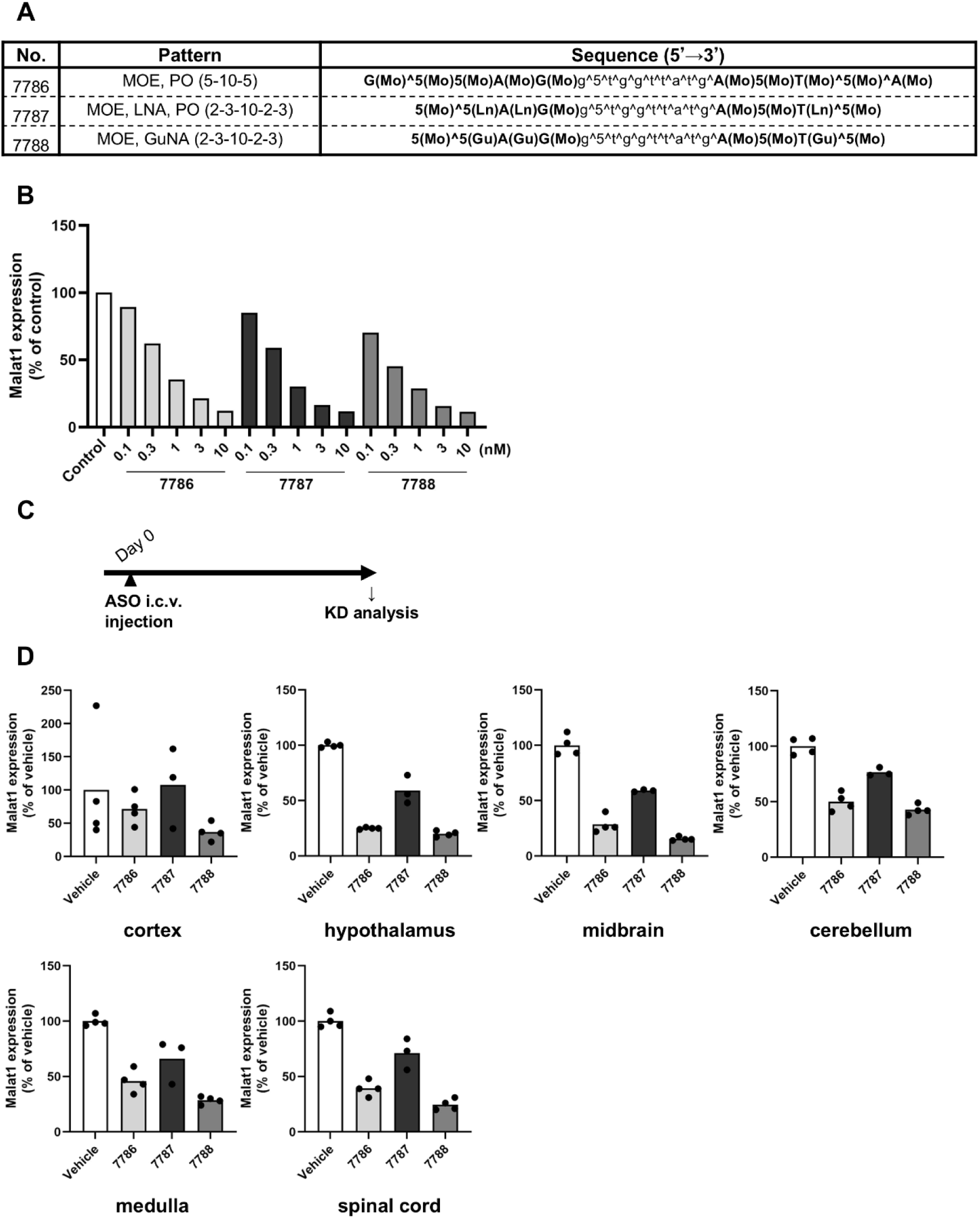
(A) Chemical modifications and sequences of modified gapmer were listed. The numbers in parentheses indicate the number of modified nucleoside in the order of 5’wing- gap-3’wing. *Gu* indicate GuNA, *Ln* LNA, *Mo* MOE, *^* phosphorothioate bond, *5* 5’-methylcytosine, *Upper letter* modified nucleoside and *lower letter* DNA. Bond between nucleoside without *^* is phosphodiester bond. (B) Percentage of mRNA expression relative to untreated control. Cells were treated with modified gapmer by lipofectamine 3000 for 24 hours. (biological replicate, n=2) (C) Schematic representation of in vivo study design and timeline. Mice were intracerebroventricular administered a single of modified gapmers at 5 nmol/head for 2 weeks. (D) Percentage of Malat1 mRNA expression in brain compartments (cortex, hypothalamus, midbrain, medulla, cerebellum and spinal cord) of mice relative to saline control. (n=4)

**Supplemental Fig.2.**
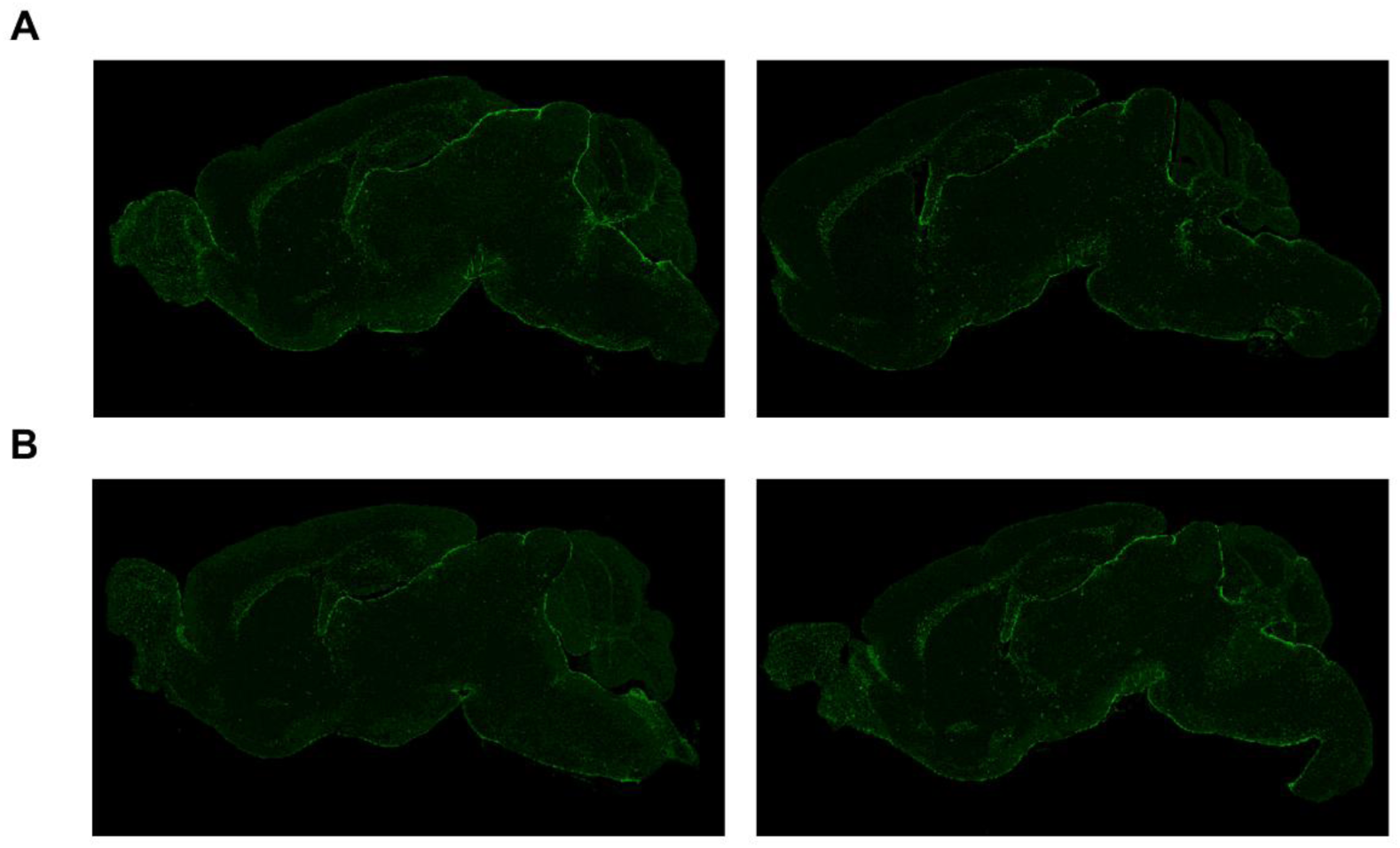
Immunohistochemistry for modified gapmer in the centrilobular area of brain from mice treated with modified gapmers or PBS and sacrificed on 2 weeks. (A) LX-A6814 (n=2), (B) LX-A8229 (n=2). Modified gapmers are shown in green.

## References

1. Scharner J, Aznarez I. Clinical Applications of Single-Stranded Oligonucleotides: Current Landscape of Approved and In-Development Therapeutics. Mol Ther 2021;29(2):540–554, doi:10.1016/j.ymthe.2020.12.022

2. Ludolph A, Wiesenfarth M. Tofersen and other antisense oligonucleotides in ALS. Ther Adv Neurol Disord 2025;18(17562864251313915, doi:10.1177/17562864251313915

3. Baek MS, Yu RZ, Gaus H, et al. In vitro metabolic stabilities and metabolism of 2’-O-(methoxyethyl) partially modified phosphorothioate antisense oligonucleotides in preincubated rat or human whole liver homogenates. Oligonucleotides 2010;20(6):309–16, doi:10.1089/oli.2010.0252

4. Khvorova A, Watts JK. The chemical evolution of oligonucleotide therapies of clinical utility. Nat Biotechnol 2017;35(3):238–248, doi:10.1038/nbt.3765

5. Geary RS, Norris D, Yu R, et al. Pharmacokinetics, biodistribution and cell uptake of antisense oligonucleotides. Adv Drug Deliv Rev 2015;87(46–51, doi:10.1016/j.addr.2015.01.008

6. Obika S, Nanbu D, Hari Y, et al. Synthesis of 2′-O,4′-C-methyleneuridine and - cytidine. Novel bicyclic nucleosides having a fixed C3, -endo sugar puckering. Tetrahedron Letters 1997;38(50):8735–8738, 10.1016/S0040-4039(97)10322-7

7. Singh SK, Koshkin AA, Wengel J, et al. LNA (locked nucleic acids): synthesis and high-affinity nucleic acid recognition. Chemical communications 1998;4):455–456

8. Kawanobe T, Asano S, Kandori H, et al. Hepatotoxicity Reduction Profiles of Antisense Oligonucleotides Containing Amido-Bridged Nucleic Acid and 2’-O,4’-C-Spirocyclopropylene Bridged Nucleic Acid. Nucleic Acid Ther 2025;35(3):114–124, doi:10.1089/nat.2024.0047

9. Sasaki T, Hirakawa Y, Yamairi F, et al. Altered Biodistribution and Hepatic Safety Profile of a Gapmer Antisense Oligonucleotide Bearing Guanidine-Bridged Nucleic Acids. Nucleic Acid Ther 2022;32(3):177–184, doi:10.1089/nat.2021.0034

10. Vester B, Wengel J. LNA (locked nucleic acid): high-affinity targeting of complementary RNA and DNA. Biochemistry 2004;43(42):13233–41, doi:10.1021/bi0485732

11. Bianchini D, Omlin A, Pezaro C, et al. First-in-human Phase I study of EZN-4176, a locked nucleic acid antisense oligonucleotide to exon 4 of the androgen receptor mRNA in patients with castration-resistant prostate cancer. Br J Cancer 2013;109(10):2579–86, doi:10.1038/bjc.2013.619

12. Swayze EE, Siwkowski AM, Wancewicz EV, et al. Antisense oligonucleotides containing locked nucleic acid improve potency but cause significant hepatotoxicity in animals. Nucleic Acids Res 2007;35(2):687–700, doi:10.1093/nar/gkl1071

13. van Meer L, Moerland M, Gallagher J, et al. Injection site reactions after subcutaneous oligonucleotide therapy. Br J Clin Pharmacol 2016;82(2):340–51, doi:10.1111/bcp.12961

14. Lovett A, Chary S, Babu S, et al. Serious Neurologic Adverse Events in Tofersen Clinical Trials for Amyotrophic Lateral Sclerosis. Muscle Nerve 2025;71(6):1006–1015, doi:10.1002/mus.28372

15. Burel SA, Machemer T, Baker BF, et al. Early-Stage Identification and Avoidance of Antisense Oligonucleotides Causing Species-Specific Inflammatory Responses in Human Volunteer Peripheral Blood Mononuclear Cells. Nucleic Acid Ther 2022;32(6):457–472, doi:10.1089/nat.2022.0033

16. Coch C, Lück C, Schwickart A, et al. A human in vitro whole blood assay to predict the systemic cytokine response to therapeutic oligonucleotides including siRNA. PLoS One 2013;8(8):e71057, doi:10.1371/journal.pone.0071057

17. Goyenvalle A, Jimenez-Mallebrera C, van Roon W, et al. Considerations in the Preclinical Assessment of the Safety of Antisense Oligonucleotides. Nucleic Acid Ther 2023;33(1):1–16, doi:10.1089/nat.2022.0061

18. Shrestha AR, Kotobuki Y, Hari Y, et al. Guanidine bridged nucleic acid (GuNA): an effect of a cationic bridged nucleic acid on DNA binding affinity. Chem Commun (Camb) 2014;50(5):575–7, doi:10.1039/c3cc46017g

19. Yamaguchi T, Horie N, Aoyama H, et al. Mechanism of the extremely high duplex-forming ability of oligonucleotides modified with N-tert-butylguanidine- or N-tert-butyl-N’-methylguanidine-bridged nucleic acids. Nucleic Acids Res 2023;51(15):7749–7761, doi:10.1093/nar/gkad608

20. Henry S, Stecker K, Brooks D, et al. Chemically modified oligonucleotides exhibit decreased immune stimulation in mice. J Pharmacol Exp Ther 2000;292(2):468–79

21. Yoshida T, Hagihara T, Uchida Y, et al. Introduction of sugar-modified nucleotides into CpG-containing antisense oligonucleotides inhibits TLR9 activation. Sci Rep 2024;14(1):11540, doi:10.1038/s41598-024-61666-3

22. Pollak AJ, Cauntay P, Machemer T, et al. Inflammatory Non-CpG Antisense Oligonucleotides Are Signaling Through TLR9 in Human Burkitt Lymphoma B Bjab Cells. Nucleic Acid Ther 2022;32(6):473–485, doi:10.1089/nat.2022.0034

23. Partridge W, Burel SA, Ferng A, et al. Correlations between preclinical BJAB assay ranking of antisense drugs and clinical trial adverse events. Clin Transl Sci 2023;16(4):575–580, doi:10.1111/cts.13476

24. Jafar-Nejad P, Powers B, Soriano A, et al. The atlas of RNase H antisense oligonucleotide distribution and activity in the CNS of rodents and non-human primates following central administration. Nucleic Acids Res 2021;49(2):657–673, doi:10.1093/nar/gkaa1235

25. Toonen LJA, Casaca-Carreira J, Pellisé-Tintoré M, et al. Intracerebroventricular Administration of a 2’-O-Methyl Phosphorothioate Antisense Oligonucleotide Results in Activation of the Innate Immune System in Mouse Brain. Nucleic Acid Ther 2018;28(2):63–73, doi:10.1089/nat.2017.0705

26. Ohto U, Ishida H, Shibata T, et al. Toll-like Receptor 9 Contains Two DNA Binding Sites that Function Cooperatively to Promote Receptor Dimerization and Activation. Immunity 2018;48(4):649–658.e4, doi:10.1016/j.immuni.2018.03.013

27. Rutz M, Metzger J, Gellert T, et al. Toll-like receptor 9 binds single-stranded CpG-DNA in a sequence- and pH-dependent manner. Eur J Immunol 2004;34(9):2541–50, doi:10.1002/eji.200425218

28. Duhamel M, Rodet F, Murgoci AN, et al. The proprotein convertase PC1/3 regulates TLR9 trafficking and the associated signaling pathways. Sci Rep 2016;6(19360, doi:10.1038/srep19360

29. Barker SJ, Thayer MB, Kim C, et al. Targeting the transferrin receptor to transport antisense oligonucleotides across the mammalian blood-brain barrier. Sci Transl Med 2024;16(760):eadi2245, doi:10.1126/scitranslmed.adi2245

30. Azevedo FA, Carvalho LR, Grinberg LT, et al. Equal numbers of neuronal and nonneuronal cells make the human brain an isometrically scaled-up primate brain. J Comp Neurol 2009;513(5):532–41, doi:10.1002/cne.21974

31. Boyd MM, Mazur C, Pribnow K, et al. Comparison of MALAT1 antisense oligonucleotide distribution following intracerebroventricular and lumbar intrathecal routes of administration. Mol Ther Nucleic Acids 2025;36(4):102742, doi:10.1016/j.omtn.2025.102742

